# Multi-State Design of Flexible Proteins Predicts Sequences Optimal for Conformational Change

**DOI:** 10.1101/741454

**Authors:** Marion Sauer, Alexander M. Sevy, James E. Crowe, Jens Meiler

## Abstract

Computational protein design of an ensemble of conformations for one protein – *i.e.*, multi-state design – determines the side chain identity by optimizing the energetic contributions of that side chain in each of the backbone conformations. Sampling the resulting large sequence-structure search space limits the number of conformations and the size of proteins in multi-state design algorithms. Here, we demonstrated that the REstrained CONvergence (RECON) algorithm can simultaneously evaluate the sequence of large proteins that undergo substantial conformational changes, such as viral surface glycoproteins. Simultaneous optimization of side chain conformations across all conformations resulted in an increase of 30% to 40% in sequence conservation when compared to single-state designs. More importantly, the sampled sequence space of RECON designs resembled the evolutionary sequence space of functional proteins. This finding was especially true for sequence positions that require substantial changes in their local environment across an ensemble of conformations. To quantify this rewiring of contacts at a certain position in sequence and structure, we introduced a new metric designated ‘contact proximity deviation’ that enumerates contact map changes. This measure allows mapping of global conformational changes into local side chain proximity adjustments, a property not captured by traditional global similarity metrics such as RMSD or local similarity metrics such as changes in φ and ψ angles.

**Author Summary:** Multi-state design can be used to engineer proteins that need to exist in multiple conformations or that bind to multiple partner molecules. In essence, multi-state design selects a compromise of protein sequences that allow for an ensemble of protein conformations, or states, associated with a particular biological function. In this paper, we used the REstrained CONvergence (RECON) algorithm with Rosetta to show that multi-state design of flexible proteins predicts sequences optimal for conformational change, mimicking mutation preferences sampled in evolution. Modeling optimal local side chain physicochemical environments within an ensemble selected significantly more native-like sequences than selections performed when all conformations states are designed independently. This outcome was particularly true for amino acids whose local side chain environment change between conformations. To quantify such contact map changes, we introduced a novel metric to show that sequence conservation is dependent on protein flexibility, *i.e*., changes in local side chain environments between stated limit the space of tolerated mutations. Additionally, such positions in sequence and structure are more likely to be energetically frustrated, at least in some states. Importantly, we showed that multi-state design over an ensemble of conformations (space) can explore evolutionary tolerated sequence space (time), thus enabling RECON to not only design proteins that require multiple states for function but also predict mutations that might be tolerated in native proteins but have not yet been explored by evolution. The latter aspect can be important to anticipate escape mutations, for example in pathogens or oncoproteins.

## Introduction

The central dogma of structural biology states that amino acid sequence defines the three-dimensional protein structure, which in turn, defines protein function, highlighting the central importance of the so-called ‘folding problem’: which three-dimensional structure adopts a given amino acid sequence? Computational protein design solves the so-called ‘inverse folding problem’ by identifying an amino acid sequence that is compatible with a given protein structure, *i.e.*, backbone conformation and possibly interactions with partner biomolecules. This approach allows for the molecule to conduct its function in this single state. Protein function, however, often relies on the transition between multiple conformations – a protein must be thermodynamically stable in multiple conformations before it is capable of achieving a defined function. Thus, for a protein to conserve its function, we hypothesized that the conservation of protein flexibility limits the protein’s sequence space to be consistent with the conformational changes needed for function.

Determining functionally relevant sequence tolerance, or rather, the set of amino acid sequences that are allowable given a protein’s function, therefore depends on identifying the set of amino acid sequences that is stable in each of the conformations needed. Testing this hypothesis is complicated, as typically not all functionally relevant conformations have been determined experimentally. The picture gets even more complicated if we look not only at functionally relevant conformations that are by definition local free energy minima (*i.e.*, thermodynamics) but also include an analysis of the height of barriers connecting these states that determine the kinetics of interconversion.

Humphris-Narayanan and colleagues demonstrated that prediction of mutation preferences of HIV-1 protease and HIV-1 reverse transcriptase was improved up to 25% when structural ensembles were included during protein design, as opposed to design of a single conformation (1, 2). Friedland and colleagues compared ubiquitin design on a single backbone conformation with design ensembles created through ROSETTA and showed that the sampled sequence space is distinct. When the ensemble conformations were selected to match experimental restraints for dynamics (nuclear magnetic resonance, or NMR, residual dipolar couplings), the sequences consistent with the ensemble match the sequences occurring in the family of ubiquitin proteins (3). These studies demonstrated that a requirement for protein flexibility substantially alters and reduces the sequence space available for evolution. One limitation of this approach, however, is that it assumes that the tolerated sequence space for a conformationally flexible protein can be determined by integrating over single-state design (SSD) of all of its conformations, *i.e.*, enumerating the most energetically favorable amino acid for each position and each conformation. However, the most energetically favorable amino acid, as determined by the aforementioned methods, may be the most energetically favorable for a single conformation, but may not be energetically tolerable at the same position in another conformation. For instance, in some conformations, the energetically most favorable amino acid might be the only allowed amino acid, with all others prohibited (imagine a tiny space where only glycine fits). At the same position in other conformations, there may be acceptable alternatives with more energetically favorable scores, but those residues could not be tolerated as an acceptable mutation in the aforementioned more constrained position. Thus, we hypothesized that multi-state design (MSD) over all conformations relevant for function will yield a more accurate representation of the biologically relevant sequence space compatible with function.

MSD approaches evaluate the stability of sequences across all conformations to select the lowest-energy sequence. The Best Max-Marginal First (BMMF) algorithm was used to demonstrate that the MSD of 16 unique calmodulin-substrate complexes increased the similarity of the designed calmodulin binding site to evolutionary sequence profiles by two-fold, and increased in native sequence recovery from 52.5% for single-state designs to 80% for the 16-state design scenario (4). Challenges for applying MSD methods like the BMMF algorithm, however, are the efficiency of the search algorithm, large memory requirements, and extended computational time needed. MSD methods up to now have been limited to designing a small number of amino acid positions across all states, with the largest number of simultaneously evaluated design positions being 27 designed positions across 60 states using the MSD FASTER algorithm (4–8).

We sought to study large proteins that undergo conformational rearrangements that include domain or hinge displacements of greater than a few Å in root mean square distance (RMSD). We expected that the tolerated sequence space must be restricted in some regions to allow for substantial ‘rewiring’ of contact networks when transitioning from one state to another. The tolerated sequence space of these types of conformational changes is not limited to local regions, such as protein-protein interfaces, but instead distributed over the entire amino acid sequence. Thus, an MSD approach that seeks to explore such sequence spaces needs to include the entire protein. The REstrained CONvergence (RECON) algorithm was used previously to estimate the sequence tolerance within protein-protein interfaces. However, already at that time this approach proved to be more computationally efficient than the generic ROSETTA MSD algorithm (8). With the addition of a message-passage interface (MPI), RECON MSD can combine the single-state design (SSD) efficiency of evaluating the sequence tolerance of a full-length protein with the MSD capability of evaluating the fitness function of a sequence across multiple conformations (9).

Highly flexible viral glycoproteins, such as the influenza A hemagglutinin protein and its stem domain (HA2), undergo conformational rearrangements of greater than 30 Å and have been shown to be conserved in sequence greater than 90% percent across subtypes (10). For other highly flexible proteins, such as calmodulin, kinases, and voltage-gated sensory channels, regions known to mediate conformational change can be conserved up to 100% across phylogenies, suggesting that a limited set of sequences is suitable for select conformation transitions (11–13). Here we use the ROSETTA RECON MSD algorithm to demonstrate that the sequence space consistent with all experimentally determined conformations of a protein approximates sequence profiles observed in evolution.

## Results

It is our hypothesis that simultaneous evaluation of sequence space across an ensemble of conformations improves the correspondence of the designed sequences to an evolutionary sequence profile, but limits the set of possible sequences by selecting mutations that are tolerated across all states within an ensemble, as opposed to individually selecting sequences that are optimal for individual states. To address this hypothesis, we perform RECON MSD and compare the designed sequences to SSD results and PSI-BLAST profiles. We quantify the similarity between designs and evolutionary sequence profiles (Fig 1A and 1B).

**Fig 1.**
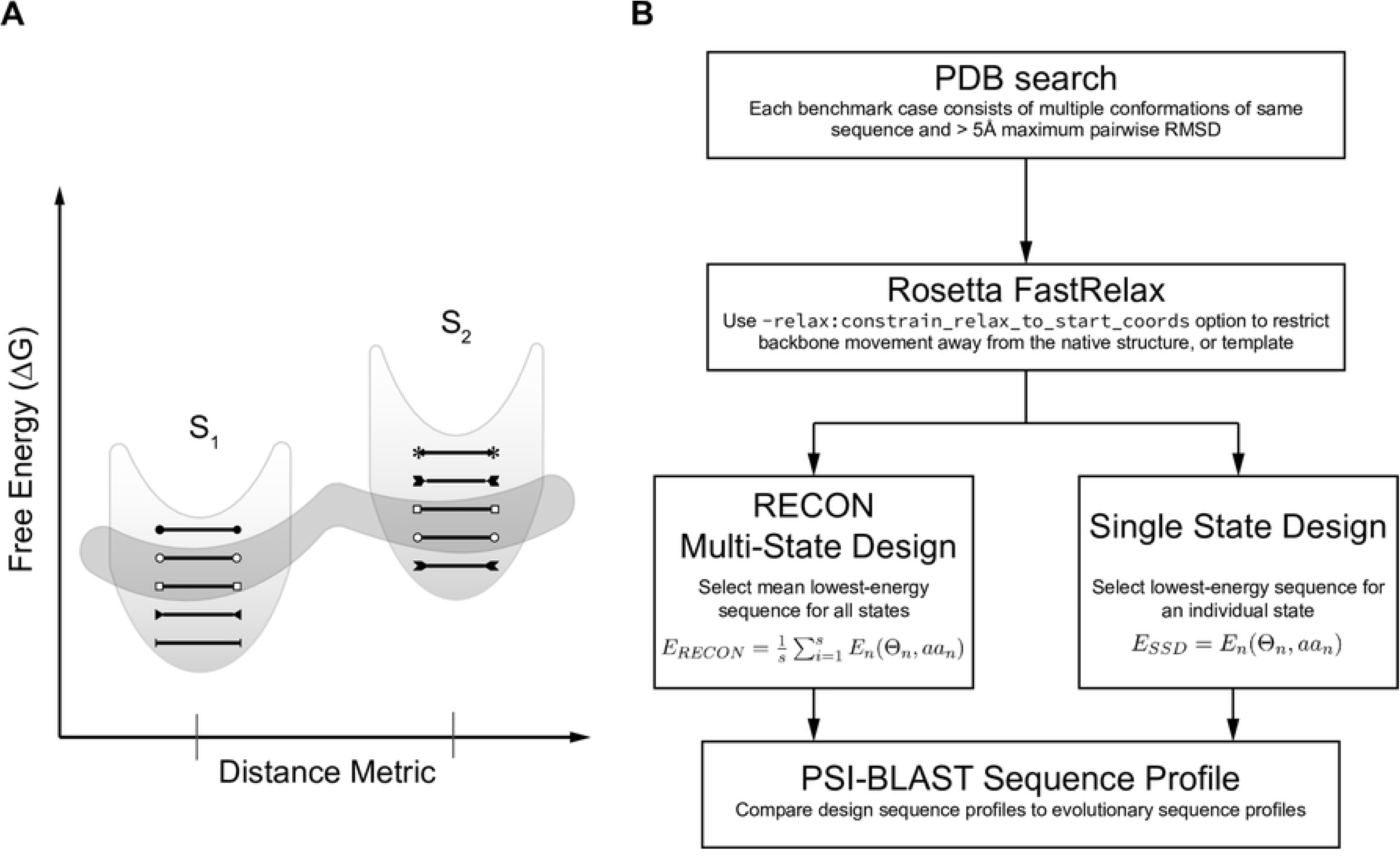
Graphical representation of hypothesis and experimental design. (A) Schematic of sequence space and the impact of flexibility on sequence tolerance. S_1_ and S_2_ represent two unique conformations of the same residue length separated by some RMSD that populate two local energy minima. Black lines with end caps represent unique sequences that are energetically most favorable for a single conformation. The dark shaded area encircles sequences that are energetically favorable for both conformations. Here we illustrate that by using multiple conformations during protein design, we identify sequences that are energetically suitable for conformational flexibility, yet are not necessarily the most stable sequence for any given conformation. Additionally, the requirement to adopt multiple conformations constrains the number of suitable sequences (B) Flow chart of benchmark design.

### Compilation of a benchmark set of eight proteins

We selected proteins with multiple known conformations of identical sequence from the PDBFlex database (14). The benchmark included eight proteins, requiring that each benchmark case have at least two published conformations with an RMSD greater than 5 Å, and an identical sequence greater than 100 amino acids in length (Table 1).We omitted duplicate conformations, which we define as conformations with and RMSD of less than 0.5 Å, to avoid bias towards similar conformations. In addition, we used a resolution cutoff of 5 Å with the requirement that greater than 75% of the included models within each design ensemble were determined at a resolution of better than 3 Å. We also omitted any models with longer sequence gaps or missing density. For structural models with chain breaks that had missing density for only one or two consecutive residues (PDB IDs 1OK8, 3C5X, and 3C6E of the dengue virus E protein monomer) we added the missing densities with the Rosetta loop modeling application (15). All structural models were gently relaxed with a restraint to start coordinates to remove any energetic frustrations frequent in models derived from low-resolution experimental structures.

**Table 1.**
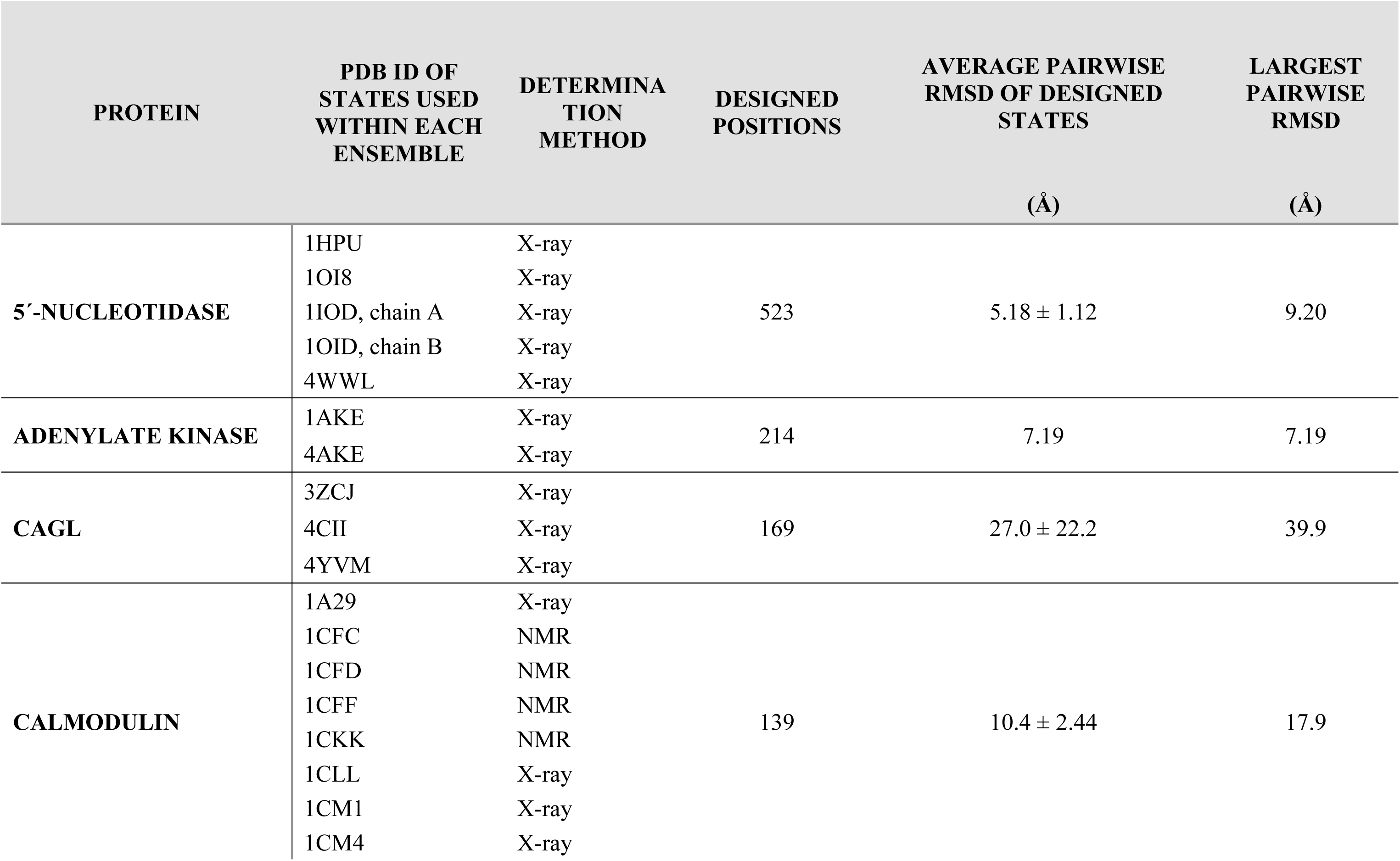
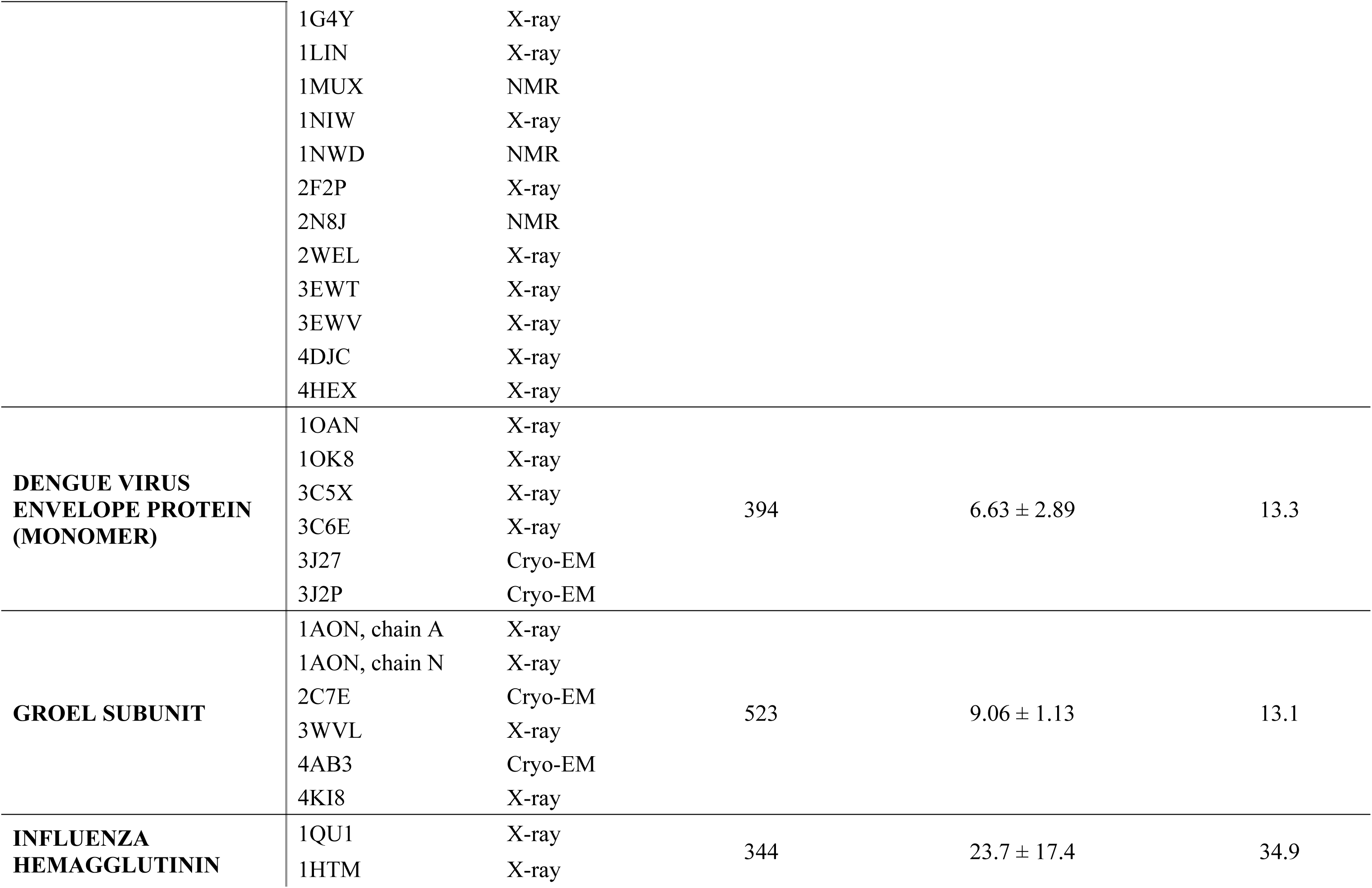
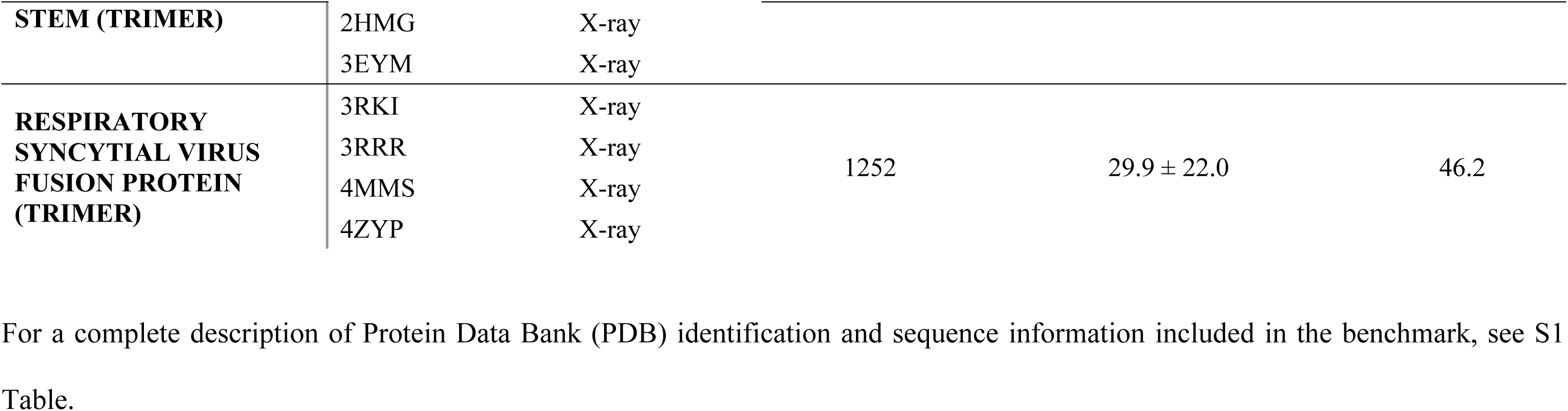
Proteins used in conformation-dependent sequence tolerance benchmark.

### Metrics to measure amplitudes of local and global conformational change

Quantification of protein flexibility commonly relies on the structural comparison of two structural models, whether that be through the similarity of equivalent atoms in three-dimensional space, calculated as root mean square distance (RMSD), or by the similarity of equivalent φ and ψ backbone dihedral angles, calculated as root mean square deviation (RMSD_da_) (16). RMSD is used frequently as a global metric used to describe the overall similarity of two conformations of the same protein and has been a powerful metric to quantify overall structural similarity. RMSD_da_, on the other hand, is used to describe local backbone displacements and is well-established, for example, to compare loop conformations. The disadvantage of both metrics is that they do not capture whether or not a particular residue is reconfigured in its interactions with neighboring amino acids. However, we hypothesize that such a metric of local rewiring driven by a global conformational space will best correlate with restrictions in sequence space introduced through conformational flexibility. Thus, we settled on three metrics that capture the structural *dis*similarity of a protein ensemble in terms of its maximum global structural dissimilarity, local backbone dissimilarity, or contact map dissimilarity: 1) The maximum pairwise RMSD of all atom coordinates of two superimposed structures within a set of *n* superimposed structures was used as a metric to describe the maximal global conformation change an ensemble undergoes (Fig 2A). To allow for comparison of RMSD values between benchmark cases that involve proteins of different size, we used RMSD100, a RMSD value normalized to protein of length 100 amino acids. (17) 2) Residue ϕ and φ RMSD_da_ was used as a local metric of similarity (Fig 2B). This metric will directly identify hinge regions between moving domains. 3) Lastly, we designed a metric that captures changes in the contact map computed as C_β_ − C_β_ distance variation. This metric captures local changes in the environment of a residue by including non-local tertiary contacts in the analysis. Thus, it is designed to capture the local and global changes of the physicochemical environment of a residue and thus defines which amino acids are tolerated in a certain position (Fig 2C and 2D). For a complete description of each metric, see Methods.

**Fig 2.**
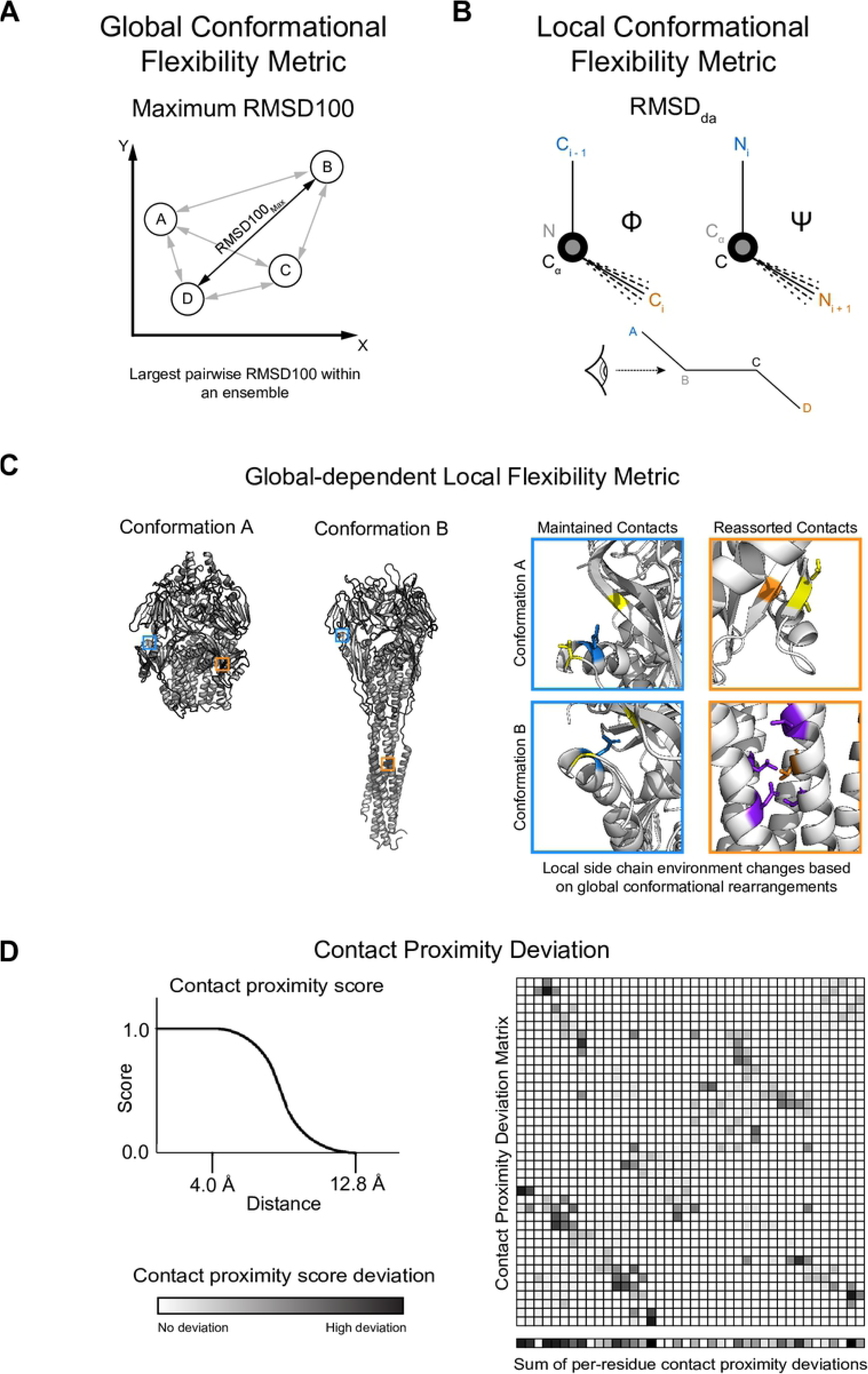
Metrics used to quantify conformational flexibility. (A) Illustration of maximum RMSD100, the metric used to quantify large-scale, or global, conformational flexibility. For simplicity, we only represent RMSD on a two-dimensional plane, where the x and y axes represent the difference in distance of cartesian space if two conformations were superimposed onto the same coordinate system. Each protein conformation of identical sequence is represented as a circle, and is separated by some distance vector evaluated as the RMSD100 of two conformations. The maximum RMSD100 describes the greatest pairwise RMSD100 within an ensemble. (B) Illustration of dihedral angle *ϕ* and *φ* variation used to calculate dihedral angle RMSD (RMSD_da_). Orientation of atoms is color-coded and corresponds to the diagram drawn at the bottom of the panel. RMSD_da_ is illustrated as the range of dotted lines, corresponding to the deviation in relative orientation of the third and fourth atoms. (C) Explanation of contact proximity deviation. Two conformations of the same protein are depicted in the left, with two residues, outlined in cyan or orange, shown in their respective positions. These two residues are magnified (top right) in their local side chain environment in Conformation A on the top and Conformation B on the bottom. Contact residues in Conformation A are colored yellow. If the same contacts are maintained in Conformation B, contact residues remain colored yellow in the bottom two boxes. If new contacts are made, contact residues are colored in purple. Even though the cyan residue changes slightly in its relative orientation between conformations, the same contacts are maintained so that the degree of conformational flexibility is relatively low in comparison to the heptad trimer refolding, and would have a low contact proximity deviation score. In contrast, the orange residue completely rearranges its local side chain contacts between conformations as a result of the large conformational rearrangement, and would have a high contact proximity deviation score. (D) Explanation of contact proximity deviation. We assigned a score to each C_β_−C_β_ distance by applying a soft-bounded, continuously differentiable function that accounts for the proximity of two side chains and approximates the likelihood of two side chains forming a contact, illustrated in the top left of Panel D. We then calculated the deviation of each C_β_−C_β_ distance across an ensemble as shown in the matrix, with low deviation scores in white and high scores in black. The contact proximity deviation score represents the sum of all C_β_−C_β_ proximity deviations a single residue undergoes within an ensemble, as shown in the bottom row separated from the matrix.

### RECON MSD samples sequence profiles that are more similar to evolutionary observed sequence profiles when compared to SSD

We first examined the correspondence of native sequence recovery determined by MSD versus SSD designed sequences with evolution conservation rates. Native sequence recovery was calculated as the mean percentage of conservation of the starting, or native, sequence for all designed positions. PSI-BLAST profiles were generated using the native sequence, and for consistency we term the percentage of the starting sequence in a PSI-BLAST profile as the percent native sequence recovery. Simultaneously sampling across multiple conformations significantly restricted sequence sampling, or in other words, was more likely to conserve the native sequence, where RECON MSD had total native sequence recovery of 87.8 ± 4.5% versus SSD with 48.9 ± 11.1% native sequence recovery (Fig 3A). In contrast, PSI-BLAST profiles had a native sequence recovery of 82.11 ± 11.2%. Qualitatively, the PSI-BLAST profiles were much more similar to the predicted sequence tolerance of RECON MSD compared to SSD, yet a Mann-Whitney U test (18) indicated a significant difference of mean native sequence recovery of either design protocol compared to PSI-BLAST sequence tolerances, with a significance of *p* = 0.0029 for RECON MSD and *p* < 0.00001 for SSD. Total sequence recovery is a coarse approximation of sequence similarity, and fails to determine if the designed sequence profiles are sampling similar mutation preferences as observed in evolution. Therefore, we calculated a total variance score of each observed position-specific mutation profile to the corresponding PSI-BLAST profile of each protein (Fig 3B and S1 Fig). We found that in seven out of eight cases, a RECON MSD mutation profile resembled its corresponding PSI-BLAST profile more closely than the SSD mutation profile.

**Fig 3.**
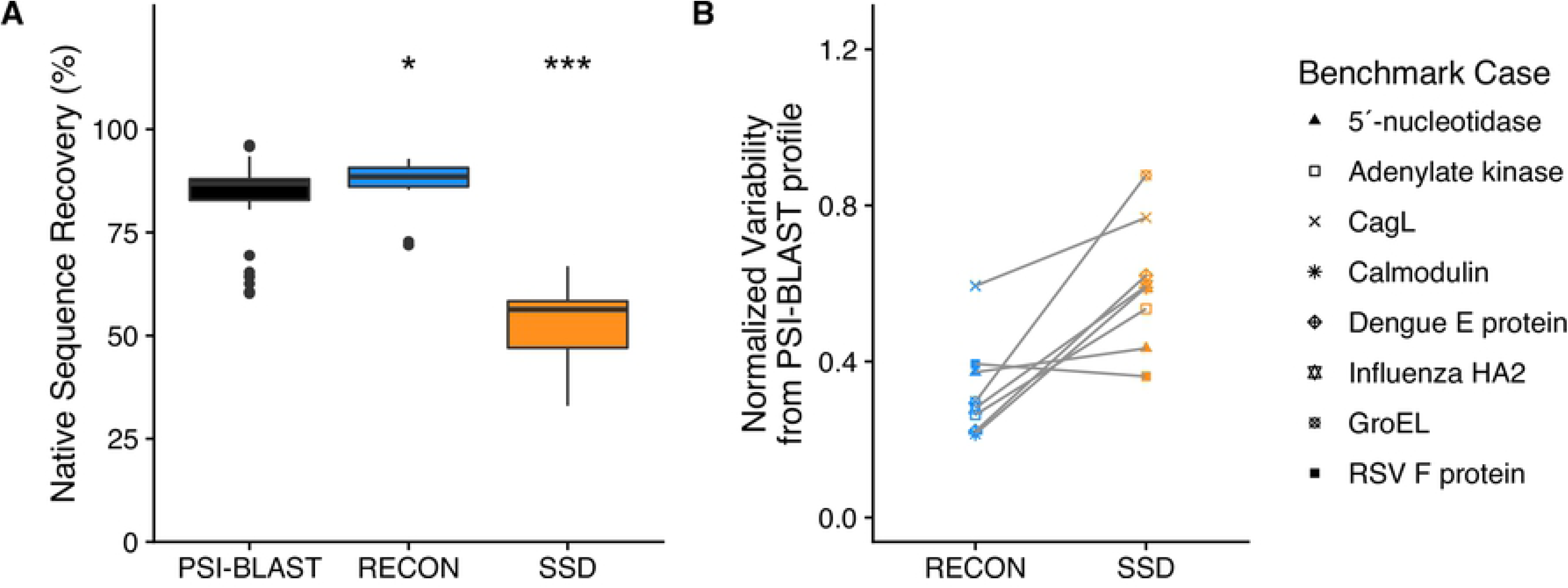
Design native sequence recovery and mutation profile variability comparisons to PSI-BLAST profiles. (A) Comparison of total native sequence recovery of relaxed and unminimized RECON MSD and SSD designs to PSI-BLAST sequence profiles generated using the native sequence. For this figure and all subsequent boxplots, shaded regions of each box plot denote values within the first and third quartiles (interquartile range, or *IQR*), with the median indicated as a solid line and whiskers representing values ± 1.5 × *IQR*. Outliers are represented as dots. Asterisks indicate the significance of difference of means of each design in comparison to the PSI-BLAST profile, with a *z*-test *p*-value < 0.01 represented by one asterisk, and a *p*-value < 0.00001 by three asterisks. The *p*-value provided in this figure and all subsequent figures represents a two-sided, 95% confidence interval. (B) Mutation frequency variances of designs in comparison to a PSI-BLAST profile, normalized by protein length. The y-axis values represent the average variability of mutation profiles for each designed residue in relation to a PSI-BLAST profile, represented as:

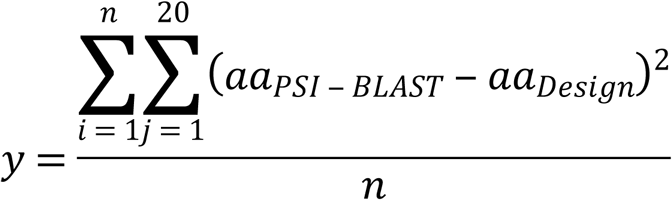

where *aa_j_* represents the frequency of an amino acid observed at position *i* for each of all twenty amino acids (*j*), and *y* is the sum of all *i* differences for all amino acids within a protein of length *n* residues. A y-value of 0 would indicate that the design profile is identical to the PSI-BLAST profile, and an increase in y-value indicates the average frequency variance of the sequence profile for each residue is more dissimilar to a PSI-BLAST profile.

### RECON MSD underestimates amino acid exchangeability, but samples a more evolutionarily relevant sequence space than SSD

Although RECON MSD more closely resembled PSI-BLAST sequence profiles on a per-case basis, we wanted to identify trends in sequence sampling in relation to the PSI-BLAST profiles to highlight design-sampling biases. This task was achieved by calculating the frequency an amino acid is conserved or mutated to another residue, or, the mean amino acid substitution frequency. In general, RECON MSD is more likely to conserve a native amino acid compared to a PSI-BLAST profile, whereas SSD is much more likely to replace the native amino acid (Fig 4A and S2 Fig). We examined amino acid exchangeability as the frequency of exchanging a native for a non-native amino acid. On average, PSI-BLAST profiles exchanged a native for non-native amino acid 1.32 ± 0.03% of the time, versus 0.77 ± 0.02% for RECON MSD and 2.45 ± 0.07% for SSD (Fig 4B). Additionally, we compared the average difference of exchangeability for each residue as observed in the PSI-BLAST profiles versus either RECON MSD or SSD and found that RECON MSD average exchangeability rates of each residue are more similar to PSI-BLAST values than SSD (Fig 4C). With the exception of phenylalanine or tyrosine, the difference between exchangeability rates for residues with larger side chains diminishes for RECON MSD, but becomes more exaggerated for SSD, as compared to observed mutation rates in evolution. This finding suggests that the inclusion of multiple conformations during design encourages better placement of bulky side chains, albeit with conservative placement. However, when comparing the linear regression model of individual exchangeability rates of either RECON MSD or SSD to PSI-BLAST rates, both designs were roughly equally dissimilar to PSI-BLAST exchangeability rates, with RECON MSD having a correlation coefficient of *r* = 0.35 and SSD with *r* = 0.64 (S3 Fig). Given that exchangeability rates were not normally distributed, a Kendall τ_*β*_ rank correlation coefficient was computed to measure the ordinal association of design and PSI-BLAST amino acid exchangeability rates, where a coefficient of τ_*β*_ = 0 would indicate that the amino acid exchangeability rates are identical. We found RECON MSD to have a τ_*β*_ = 0.283, *p* ≤ 2.22 × 10 ^‒ 16^ versus τ_*β*_ = 0.372, *p* ≤ 2.22 × 10 ^‒ 16^ for SSD when measured for its association to PSI-BLAST amino acid exchangeability rates. In addition, we compared the difference in exchangeability rates between design and PSI-BLAST by calculating the ratio of transformed exchangeability rates (*e*) as 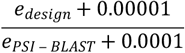 to avoid division by zero, where an individual exchangeability rate would be equivalent between design and PSI-BLAST if *e*_*ratio*_ = 1. We found that for RECON MSD the mean *e*_*ratio*_ = 2.49 and for SSD the mean *e*_*ratio*_ = 22.7. A Mann-Whitney U test of matched individual ratios found a significant difference between RECON MSD and SSD exchangeability rate ratios to PSI-BLAST exchangeability rates, with *p* < 0.0001. Taken together, RECON MSD is sampling individual mutation preferences significantly more closely to that observed in evolution than SSD.

**Fig 4.**
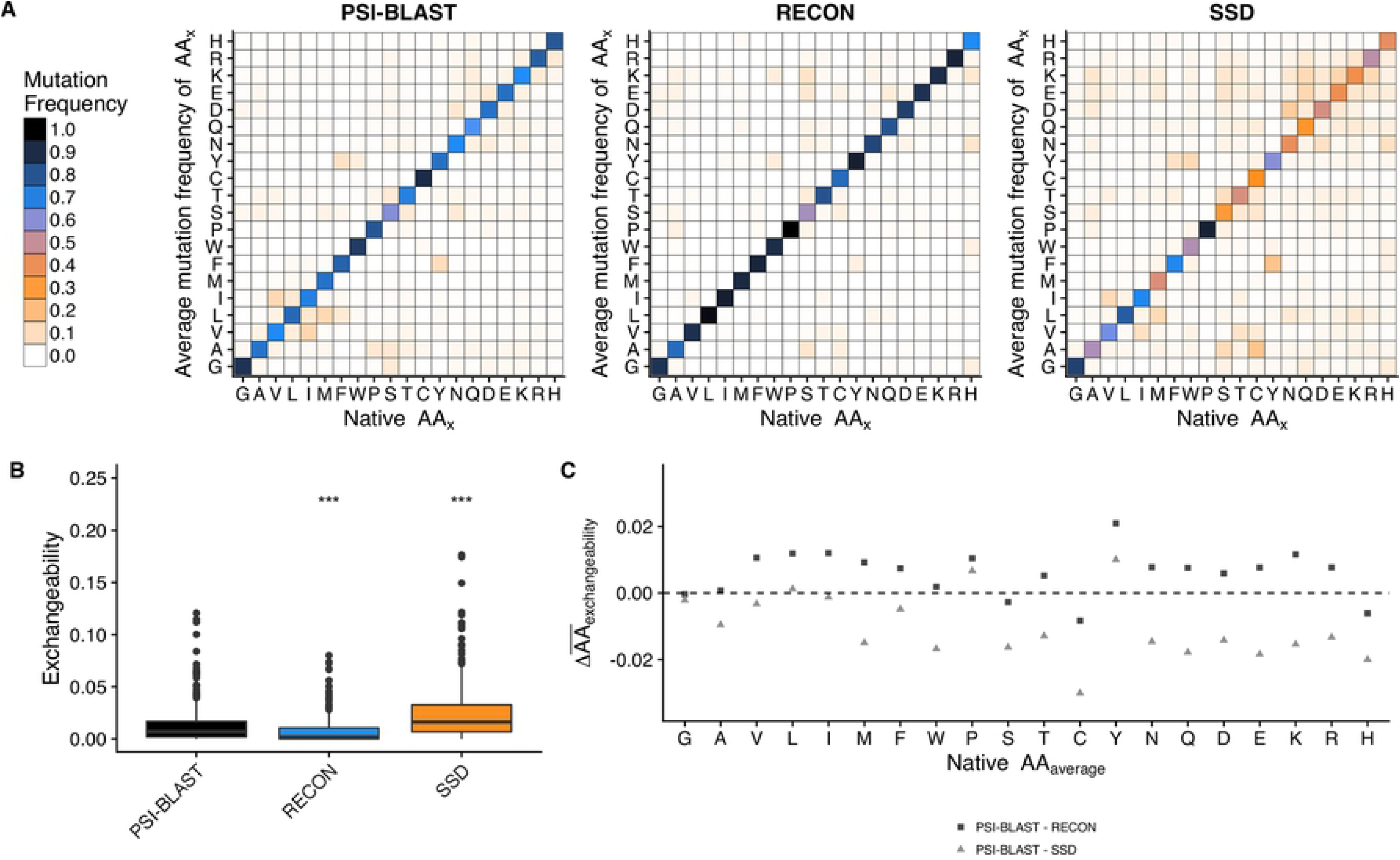
Comparison of exchangeability rates. (A) Average amino acid exchangeability of PSI-BLAST, RECON MSD, and SSD sequence profiles. Single-letter amino acid codes were used for both x and y axes, with the x axis representing the original amino acid and the y axis representing the average mutation frequency the original amino acid to the indicated mutation. (B) Comparison of exchangeability rates between profiles, excluding rates of native sequence conservation rates. The y axis represents the mean frequency a native amino acid is replaced with a specific, non-native amino acid, which we term as amino acid exchangeability. (C) Difference of mean amino acid-specific exchangeability observed in a PSI-BLAST profile compared to a design profile. The x axis represents each type of amino acid present in the native sequence. The y axis represents the difference in average exchangeability frequency of each amino acid type, or rather, the average frequency a native amino acid type is replaced with any other non-native amino acid. A positive value indicates the native amino acid is less likely to be exchanged for a non-native amino acid during design, whereas a negative value indicates the native amino acid is more likely to be exchanged, as compared to a PSI-BLAST profile.

### Sequence conservation is dependent on its contact map as computed by Cβ−Cβ distance deviations

To consider the effect of conformational flexibility on sequence conservation, we examined the dependency of native sequence recovery on different aspects of conformational flexibility using the aforementioned metrics, maximum RMSD100, RMSD_da_, and contact proximity deviation. We performed a Kendall τ_*β*_ rank correlation test on each profile to test for the strength of dependency of native sequence recovery on each metric (Fig 5). Of the three metrics, the native sequence recovery, or rather percent conservation, observed in PSI-BLAST profiles was only dependent on contact proximity deviation *z*-score, with *p* = 1.79 × 10 ^‒ 6^, versus *p* ≥ 0.144 for all other tests. RECON MSD and SSD native sequence recovery depended on both RMSD_da_ and contact proximity deviation *z*-score (*p* < 0.01). Native sequence recoveries of both designed profiles depended strongly on dihedral angle deviation, with *p* ≤ 2.22 × 10 ^‒ 16^, and had similar τ_*β*_ coefficients, with τ_*β*_ = 0.101 for RECON MSD and τ_*β*_ = 0.0806 for SSD. This finding suggests that the Rosetta scoring function employed by both protein design algorithms is too restrictive in sampling for residues at hinge points, given that the same dependency on RMSD_da_ is not observed for PSI-BLAST sequence conservation. However, both PSI-BLAST and RECON MSD had similar τ_*β*_ coefficients predicted with the same confidence, with τ_*β*_ = 0.0639, *p* = 1.79 × 10 ^‒ 6^ and τ_*β*_ = 0.0787, *p* = 1.19 × 10 ^‒ 7^ respectively, for the dependence of native sequence recovery on contact proximity deviation *z*-score. This observation suggests that there is an evolutionary constraint on residues that are required to maintain a re-assortment of their local physicochemical environments necessary for a conformational change, and that RECON MSD closely models this evolutionary constraint by considering the multiple local side-chain environments within a protein ensemble.

**Fig 5.**
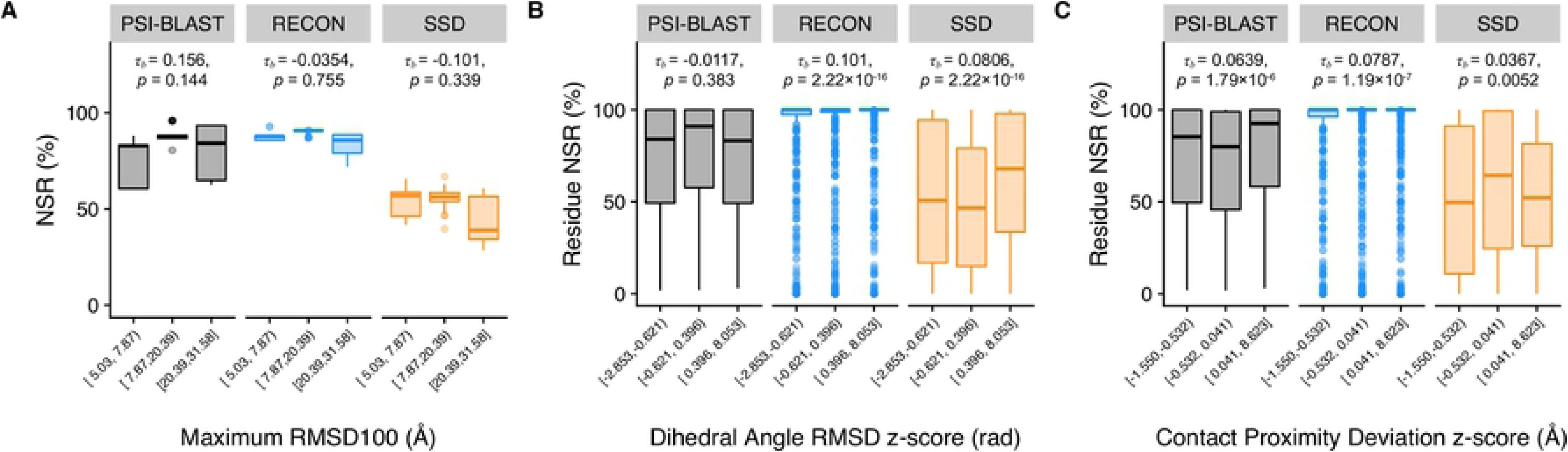
Relationship of conformational flexibility and native sequence recovery by sequence profiles. The x axis is binned into three groups of equal number of data points to show the distribution of native sequence recovery between groups of low, middle, and high values for each metric. A Kendall τ_*β*_ rank correlation test was performed on each profile to measure the strength of dependence of native sequence recovery on each metric, indicated in each plot along with its associated *p*-value. (A) Comparison of native sequence recovery dependence on maximum RMSD100 between sequence profiles. (B) Comparison of native sequence recovery dependence on RMSD_da_ between sequence profiles. RMSD_da_ values of each protein were not equally distributed, nor were of similar range. Therefore, a *z*-score of was used to normalize RMSD_da_ values of each protein to compare dihedral angle deviation scores, shown along the x axis. A similar approach was implemented to normalize contact map deviation scores. (C) Comparison of native sequence recovery dependence on contact deviation scores.

### Regions critical for conformational plasticity are energetically frustrated

The encouraged sequence convergence employed by the RECON MSD algorithm identifies amino acid sequences that have the lowest total energy across all states (8). To examine the energetic impact of requiring a single amino acid sequence to adopt multiple states, we use a similar energy score term described previously as the sum total energy score normalized by the number of designed positions (see Methods). For RECON MSD designs, this approach would include lowest mean energy score of the designed ensemble, whereas the SSD energy score would include the lowest energy scores for each state. In all eight cases, RECON MSD selects sequences with a significantly higher energy score than SSD with a paired student’s t-test (19), with *p* < 1 × 10 ^‒ 4^. (Fig 6). We also compared the design energy scores to the ten lowest-energy relaxed structures, which only included the native sequences, and found that RECON MSD samples a lower energy sequences than the relaxed native structures. Even though SSD sequences are the most stable, RECON MSD identifies more favorable mutations within the native ensemble of states.

**Fig 6.**
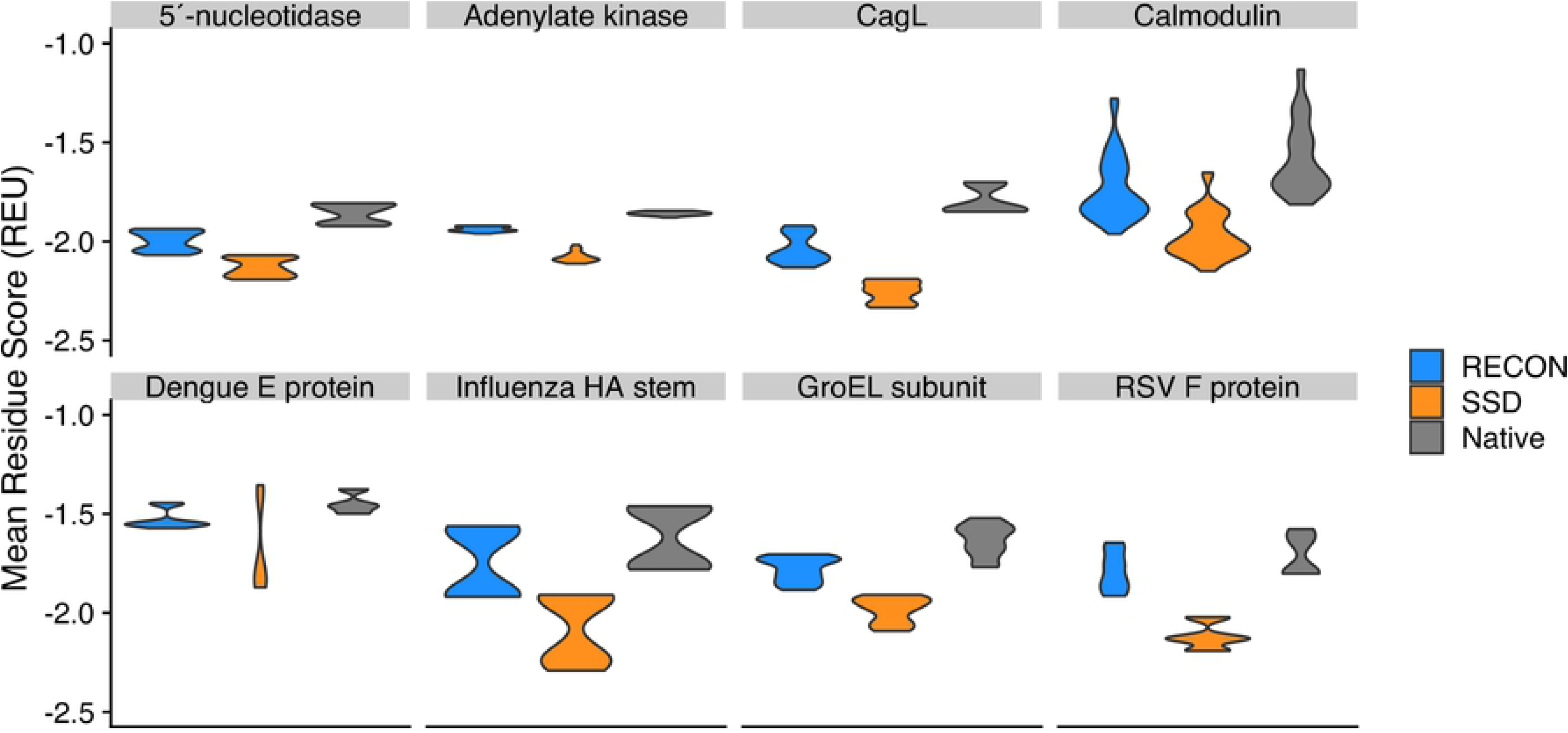
Average per-residue total energy score of the lowest ten percent scoring models for RECON MSD, SSD, and starting relaxed (Native) models. One hundred simulations were performed for each group and the lowest ten total energy scoring models were used for the comparison. The total scores were normalized so that the calculated total score was divided by the number of residues within each model to obtain a mean residue score. For RECON MSD models, the total calculated score also had to be normalized by the number of states within each model. The violin plot width indicates the normalized energy score density of each group.

### Stability decreases for residues with larger Cβ−Cβ contact map deviations

We used a Kendall τ_*β*_ rank correlation test to analyze the dependency of the modeled sequence energy score on global and local conformational changes. For the comparison with global conformational changes, we compared the mean total score of the ten lowest-energy scoring design models, normalized by the number of residues within each protein, to the maximum RMSD100 of an ensemble. We found that there is a negative dependence of mean total score on the maximum RMSD100 for SSD models (τ_*β*_ =‒ 0.143, *p* = 1.16 × 10 ^‒ 5^), but not so for RECON MSD models (τ_*β*_ = 0.0177, *p* = 0.586; **Error! Reference source not found.**). Conversely, there was a small, but significant positive dependence of individual residue scores on contact proximity deviation *z*-scores for RECON MSD models (τ_*β*_ = 0.0356, *p* = 0.00584), but not so SSD models (τ_*β*_ =‒ 0.00538, *p* = 0.677). There was no dependence of individual residue scores on dihedral RMSD for either design approach (Fig 7). This finding suggests that RECON MSD is restricted in optimizing the stability of residues that must rearrange their local side-chain environments, but not for optimizing local backbone flexibility. SSD, on the other hand, is not restricted in optimizing side-chain placement within an ensemble, and therefore can select amino acid sequences that are more stabilizing for individual conformations.

**Fig 7.**
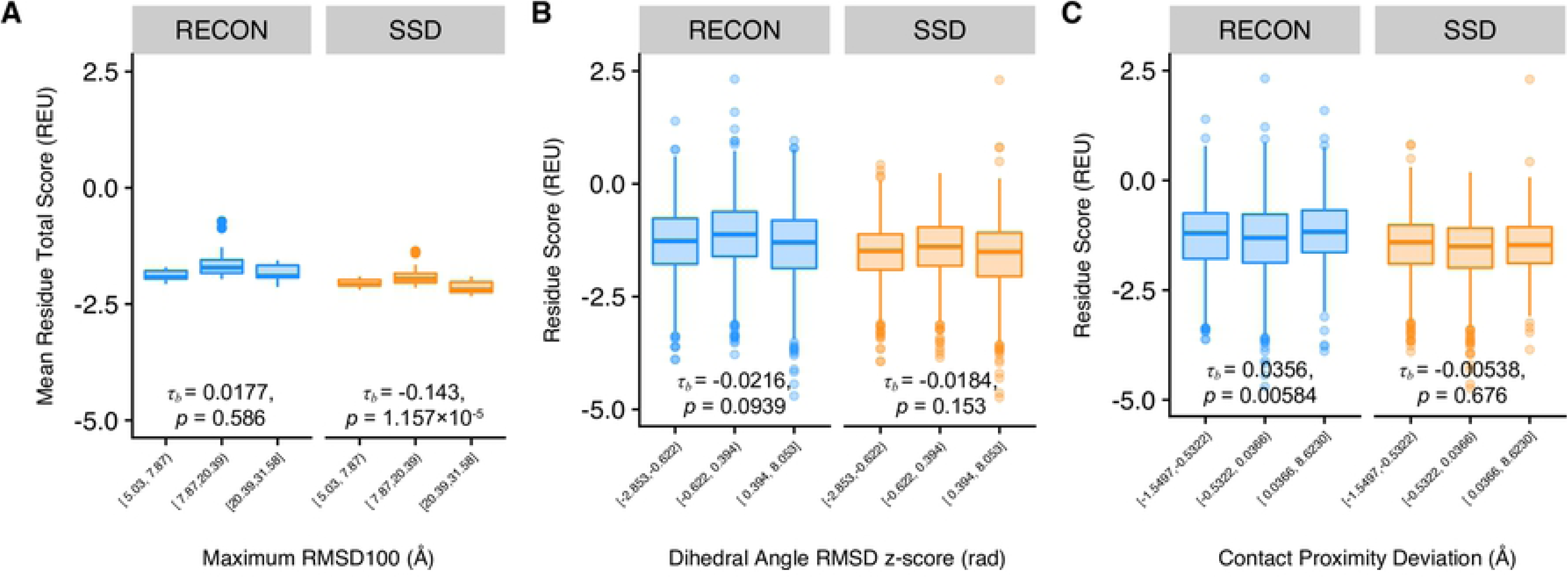
Comparison of conformational diversity and per-residue total scores. All panels are binned into low, medium, and high x values, with equal number of data points for each bin. A Kendall τ_*β*_ rank correlation test was performed on each profile to measure the strength of dependence of native sequence recovery on the x axis value, indicated in each plot along with its associated *p*-value. (A) Comparison of maximum RMSD100 and mean total energy score, normalized by the number of residues. (B) Comparison of normalized RMSD_da_ *z*-score and mean total energy score of each residue. (C) Comparison of normalized contact proximity deviation *z*-score and mean total energy score of each residue.

## Discussion

### Contact proximity deviation captures local and global conformational rearrangements as a single metric

Methods like Local-Gobal Alignment and contact area differences are useful in circumventing the over-estimation of global structural dissimilarity by either emphasizing local backbone segment structure similarity or by side chain placement similarity, respectively (20, 21). In particular, contact area differences between two homologues have been shown to be as accurate as RMSD, if not more, in comparing the structural similarity of proteins with very high sequence similarity (22). However, contact area differences are sensitive to errors in side chain atom placement within a structural model, so that accounting for side chain mutations while measuring the relative position of equivalent residues is not feasible.

For this paper, we introduced the contact proximity deviation metric that quantifies the relative placement of a side chain within a structure and its variability within an ensemble without having to predict side-chain contacts. The soft-bounded function we use to score the distance of a side chain C_β_ atom to all other surrounding side chains is less sensitive to the misplaced assignment of atom coordinates in an individual structure. Additionally, because we use the deviation of contact proximity within an ensemble rather than the contact proximity score of an individual C_β_ − C_β_ distance, small errors in the C_β_ atom assignment of one structural model are less likely to substantially influence the score deviation of a residue within an ensemble. Therefore, scoring C_β_ − C_β_ distance deviations of each residue allows for the comparison of C_β_ placement and local protein fold variability within an ensemble, independent of side chain identity.

The measurement of local structural variability by NMR residual dipolar coupling has shown that regions of high flexibility within ubiquitin ensembles align with multiple protein-protein interfaces (23). Large protein dynamics are not suitable for NMR and are difficult to experimentally determine as an ensemble. In the few cases where multiple conformations of large proteins have been resolved, contact proximity deviation provides a unique method to explore how side chain interaction flexibility or protein re-folding affects binding interfaces. Additionally, in cases such as protein switch design or structural analysis of mutations where protein flexibility, but not sequence, needs to be conserved, it is necessary to have a metric that allows for the comparison of local structural variability due to protein refolding, such as contact proximity deviation.

### Sampling functional mutation preferences requires evaluation of sequence stability as an ensemble

MSD approaches use the energetic contributions of multiple conformations to steer sequence selection and cull any sequences that do not improve the energy score of the designed ensemble as a whole (24). For most approaches, sequences that do not improve all or the majority of conformations within an ensemble are culled, which is appropriate when the goal is to stabilize a protein ensemble within an energy minimum. However, protein function may not select for sequences that are limited to a single energy minimum. To better estimate the mutational preferences critical for function, it is necessary to use an approach that models local side-chain environments within the context of an ensemble.

Within Fig 4, we illustrated that, although RECON MSD failed to accurately predict all amino acid exchangeability rates, the consideration of multiple local side chain environments during protein design improved the prediction of sequence conservation and overall accuracy of amino acid substitution frequencies as compared to modeling local side chain environments independently, particularly when modeling bulky side chains. In combination with Fig 6 and Fig 7, we demonstrated that the consideration of local side chain stability within the context of an ensemble restricts stability optimization of the ensemble, especially for residues that require side-chain rearrangements during a conformational rearrangement. Taken together, we demonstrated that selection of mutation profiles by RECON MSD is much more similar to mutation rates observed in homologs if each mutation is evaluated across every conformation, or state, within an ensemble, and then culled if the mutation is evaluated to be destabilizing for an individual state within an ensemble. This approach does not necessarily select sequences that improve the stability of every state within an ensemble, but rather places the importance of modeling an ideal conformation-specific, local side-chain environment to prevent local side-chain destabilization within the context of an ensemble.

### RECON MSD can be used to predict evolutionary sequence conservation of flexible proteins

Sequence similarity searches, such as PSI-BLAST, are fast and easy to use. Predicting mutation preferences from structure, especially if the sequence is known to form multiple conformations, remains to be a challenge. Advances in structure-based evolution design methods rely on iterative approaches that match sequence and structure similarities to predict sequence entropy (25). For proteins that undergo conformational rearrangements, using this type of approach to search for structural similarity limits the sequence search space to similar conformations, possibly preventing the identification of sequences capable of adopting multiple conformations. Although other MSD methods have improved the selection of more evolutionarily similar sequences, they are limited in their capacity to simultaneously sample conformation and sequence space so that the relevance of conformational plasticity in evolutionary dynamics have not been fully interrogated.

For this benchmark, we used RECON MSD to demonstrate that sequence conservation and mutation preferences of a single sequence can be approximated using the evaluation of local residue physicochemical changes, provided that this one sequence folds into select, multiple conformations. In Fig 3, we showed that the estimated sequence conservation of RECON MSD designs differs by roughly 5% from the sequence conservation observed in PSI-BLAST profiles, with RECON MSD being more conservative. We demonstrated in Fig 5 that sequence conservation, as observed in evolutionarily-related sequences, depends on the degree of local side-chain environment changes a residue must undergo to assist in the refolding of a protein during a conformation rearrangement, and that RECON MSD approximates the same restriction in amino acid conservation as C_β_−C_β_ proximity variation increases.

The caveat to using RECON MSD to predict mutation preferences is accounting for ROSETTA sampling biases. First, RECON MSD does not currently allow for the formation or destruction of disulfide bonds, which is critical for conformation stability, and does not accurately model the frequency of cysteine conservation. Consideration of alternate protonation states due changes in pH are also not explicitly modeled, which we see from our amino acid exchangeability comparisons that RECON MSD underrepresents exchangeability of polar residues and frequently mutates histidine to lysine or arginine, which has a pK_a_ much higher than histidine or which is not charged. Additionally, in Fig 5, we showed that RECON MSD is likely to overestimate sequence conservation of hinge regions that have large dihedral angle RMSDs. Even though we used a gentle minimization prior to design, minimization significantly increases the estimated stability of the native residue, making the replacement of the native amino acid unfavorable, as shown in S1 Fig. Given that residues located at hinge points within flexible loops are intrinsically disordered and typically contain less than ideal Ramachandran dihedral angles, it is likely that minimization specifically overcorrects these bond angles to fit the energy scoring function, preventing accurate sampling of rotamer placement. With the addition of explicit disulfide bond formation, use of a pK_a_-dependent rotamer library, and improvement of minimization prior to design, the RECON MSD algorithm could prove to be a valuable tool in predicting accurate mutation profiles.

The computational time required for the RECON MSD design simulations within this benchmark ranged from 2 - 36 hours. Compared to experimental approaches that have tested for functionally tolerated mutations in either dengue virus envelope protein or influenza hemagglutinin protein (2, 26, 27), RECON MSD is much faster and less costly in identifying biologically relevant mutations. Additionally, RECON MSD is not limited to sampling mutations singly, pairwise, or as limited networks, but rather can sample mutations as an interaction network of each local side-chain environment. Traditional intra-protein co-evolution methods, such as direct coupling analysis (28), mutual information (29–31), or McLachlan-based substitution correlation methods (32, 33), are not reliable in detecting co-variation or correlation of mutation frequencies of highly conserved sequences (34), and so they fail to detect contact dependencies of sequences with low sequence variation. In the case of this benchmark, we see that flexible sequences tend to be more highly conserved, especially when residues need to maintain distinct contacts between conformations. Therefore, current co-evolution methods cannot be used to detect residue contact dependencies of flexible, highly conserved sequences, whereas this benchmark suggests that RECON MSD is well-suited to identifying the evolutionary potential of a flexible sequence.

## Conclusions

We demonstrated that RECON MSD significantly improves the similarity to evolutionary mutation preferences from SSD selected mutation profiles by selecting sequences which are energetically favorable for an ensemble of local side-chain interactions. Specifically, in instances where the goal of protein design is to preserve an ensemble of conformations for functionality, we suggest a greater emphasis on designing local physicochemical environments for each and all conformations within an ensemble, and to place less of an emphasis of finding sequences representing the most thermostabilizing for either each state individually or as an average of all states. Furthermore, the new conformational diversity metric contact proximity deviation we describe in this paper allows for the comparison of protein ensembles, assuming they are of similar length but not sequence, by quantifying position-specific relocation due to one or more conformational changes. Therefore, in conjunction with contact proximity deviation, RECON MSD warrants further use as a bioinformatic tool to estimate mutation preferences of homologous proteins, especially for proteins known to undergo similar domain or fold reorganization between conformations.

## Methods

### Selection and preparation of benchmark datasets

Our criteria for benchmark datasets included proteins that had at least two published conformations with greater than 5 Å RMSD and at least one peptide chain greater than 100 residues in length. To identify these proteins, we performed a BLAST search to identify proteins with 100% sequence identity and with gaps of three or less residues in length. Structures with similar backbone conformations of less than 1 Å RMSD were excluded from design so that the structure with the longest matching consecutive sequence was kept as the template structure.

Structures were downloaded from the Protein Data Bank (PDB; www.rcsb.org) and processed manually to remove all atoms other than the residue atoms intended for design. Any residues that did not align or positions that were not present in all template structures were not considered for design and were removed from the template. For a detailed description of which residues were included for design, see S1 Table. Template structures were subject to minimization and repacking in ROSETTA using FastRelax constrained to start coordinates with a standard deviation of ± 0.5 Å with the talaris 2013 score function (35, 36). The lowest total energy score model of 100 relaxed models was selected for design. For comparisons using un-relaxed models, the template structure was the input model structure used for relaxation.

### RECON MSD multi-specificity and single-state design

Benchmarking using RECON MSD multi-specificity design was performed using four rounds of fixed backbone design and a convergence step using the greedy selection algorithm, as previously described (8), with the exception that only repacking, and not backbone minimization was allowed following the convergence step to prevent over-optimization of the energy score following design. Similarly, single-state design was performed using four rounds of fixed backbone rotamer optimization followed by repacking using the identical designable residues as specified for RECON MSD designs. The talaris 2013 scoring function was used for both RECON MSD and single-state designs. For each protein dataset, 100 designs were generated for each benchmark structure using either RECON MSD or SSD.

### Generation of sequence profiles

The lowest ten out of a hundred scoring models were used for quantification of sequence tolerance. In the case of SSD, the ten lowest total scoring models were used from the design simulation of each PDB structure. For RECON MSD, the total score of each model designed within an ensemble design run were added to create a fitness score, which then was sorted to identify the ten designed ensembles with the lowest fitness score. A Shannon entropy bitscore was calculated for each designed position within an ensemble as *I*_*i*_ = *p*_*i*_ × log_2_ (20 × *p*_*i*_), with *i* as the amino acid and *p*_*i*_ as the frequency of that amino acid. Here, the calculated amino acid frequency includes the frequency at the same position within the ten lowest-scoring models of all designed states, whether designed independently by SSD or designed simultaneously by RECON MSD, such that an amino acid represented a 100% of the time at a particular position in all states has a bitscore of 4.32 (37).

PSI-BLAST profiles were obtained by querying a non-redundant protein database using default parameters, increasing the number of iterations to ten iterations, as well as querying the database with *e*-value thresholds ranging from 1 × 10 ^‒ 5^ to 1 × 10^2^. We reported only PSI-BLAST profiles generated using default parameters, which includes two iterations and an *e*-value of 0.005. Comparisons to PSI-BLAST profiles using non-default parameters were qualitatively identical to the PSI-BLAST profiles generated using the default parameters, and were therefore not reported. We omitted any sequences within the queried sequence profiles which were not included for design and calculated the Shannon entropy bitscore for each position using the position specific-scoring matrix (PSSM) frequencies as described in the previous paragraph.

### Comparison of sequence profiles

We compared the position specific-scoring matrix (PSSM) generated from the PSI-BLAST query to the PSSM generated by either RECON MSD or SSD by calculating the percentage of native sequence recovery, which was determined as the sum of the bitscores of the native amino acids at each position divided by the sum of the information bitscore of all amino acids at all positions (38). Additionally, we calculated the difference in designed position mutation frequencies from each position with the PSSM, as described in **Error! Reference source not found.**:

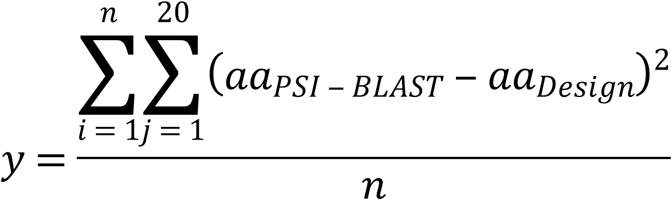

where *aa_j_* represents the frequency of an amino acid observed at position *i* for each of all twenty amino acids (*j*), and *y* is the sum of all *i* differences for all amino acids within a protein with a length of *n* residues. Average amino acid substitution rates were determined as the mean cumulative substitution frequency of each amino acid *i* to amino acid *j*, where

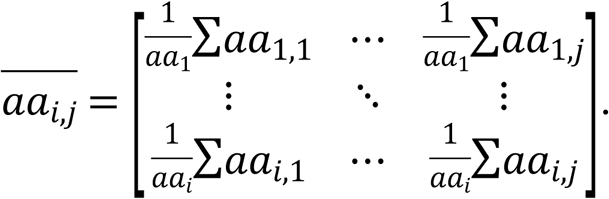

We define amino acid exchangeability as the subset of average amino acid substation rates that exclude the substitution rates of amino acid *i* to amino acid *j*, where *j* is identical to *i*, or in other words, all substitution rates that include the average conservation frequencies of the native amino acid. The mean amino acid exchangeability rates, as shown in **Error! Reference source not found.**C, were calculated as

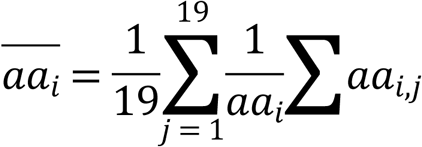

where the mean exchangeability rate of each amino acid *i* is the mean of all exchangeability rates of amino acid *i* to amino acid *j*, excluding the conservation rate of amino acid *i*. Kendall τ_*β*_rank comparison tests, linear regression, Wilcox comparison of means, and student t tests were performed in R.

### Description of conformational metrics used in this benchmark

We use the maximum RMSD within an ensemble to represent the largest amplitude of dissimilarity within an ensemble, defined as:

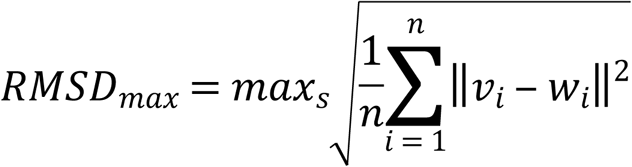

where *n* represents the number of residues, *s* represents the number of structures within an ensemble, and 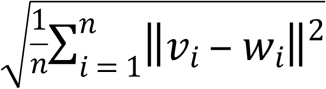 represents each pairwise RMSD within an ensemble (**Error! Reference source not found.**A). For the local backbone dissimilarity metric, we use dihedral angle RMSD to describe the deviation of each equivalent dihedral angle, or pair *ϕ* and *φ* angles, within an ensemble containing *s* structures, as

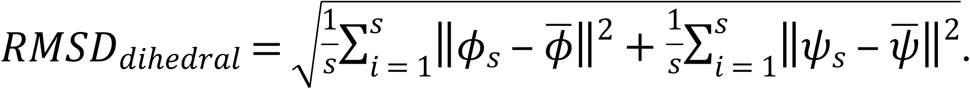

where 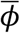 and 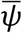 represent the mean *ϕ* and *ψ* angle of each equivalent residue. The contact map dissimilarity metric we introduce here is based off the Durham et al. (39) neighbor count weight metric, which scores the likelihood of a neighboring contact by assigning each C_β_−C_β_ distance a score, which we term contact proximity (𝒞) as

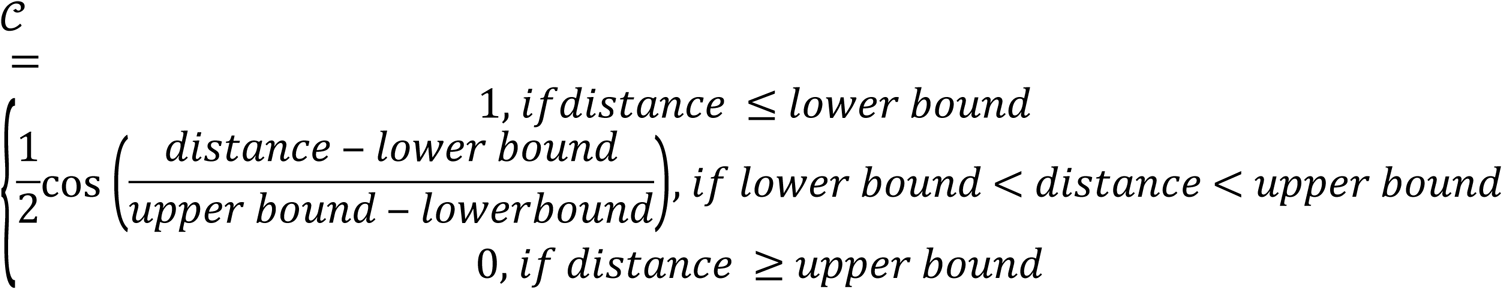

For glycines, a pseudo-C_β_ atom is defined using the amide N, C_α_, and carboxyl C atom coordinates in the PDBtools package (github.com/harmslab/pdbtools) before calculating C_β_−C_β_ distances. The lower and upper bounds represent thresholds where a C_β_−C_β_ distance certainly does and does not contain any side-chain atoms that are in contact with another residue’s side-chain atom. We define the lower bound as 4.0 Å and the upper bound 12.8 Å, where the lower bound was determined to be a reliable threshold to define solvent-inaccessible side-chains due to side-chain contacts, or in other words, a C_β_−C_β_ distance less than the lower bound is very likely to form at least one side chain interaction (39). The upper bound was determined by the maximum C_β_−C_β_ distance where at least one atom from each side chain formed an interaction (40). Finally, the contact proximity deviation for each residue was calculated as the sum of all C_β_−C_β_ contact proximity score deviations for that residue (Fig 2D). With this metric, we can quantify the changes in side chain local environments that are not due to local hinge-points, but instead, show local side chain environment changes that are due to larger conformational rearrangements.

## Acknowledgements

The authors would like to thank Amanda Duran, Axel Fischer, and Diego del Alamo for their assistance and helpful discussions during the development of this project. Calculations for the contact map deviations were made possible by members of the Michael Harms Laboratory as contributors to the PDBtools opensource software (github.com/harmslab/pdbtools). We also thank the Vanderbilt University Advanced Computing Center for Research and Education for the computational resources necessary to complete this project. All work was funded by the U19 AI117905 grant for Structure Based Design of Antibodies.

## Supplementary Information

**S1 Fig. Design native sequence recovery and mutation profile variability comparisons to PSI-BLAST profiles using relaxed and unminimized starting models.** (A) Comparison of total native sequence recovery of relaxed and unminimized RECON MSD and SSD designs to PSI-BLAST sequence profiles generated using the native sequence. Asterisks indicate the significance of difference of means of each design in comparison to the PSI-BLAST profile, with a *z*-test *p*-value < 0.01 represented by one asterisk, and a *p*-value < 0.00001 by three asterisks. (B) Mutation frequency variances of designs in comparison to a PSI-BLAST profile, normalized by protein length. The y-axis values represent the average variability of mutation profiles for each designed residue in relation to a PSI-BLAST profile, as described in Fig 3.

**S2 Fig. Difference in average amino acid exchangeability between sequence profiles.** The x axis represents the original amino acid. The y axis represents the difference in average mutation frequencies between two profiles, which are noted above each grid. Along the diagonal axis, indicating native sequence conservation, values less than zero (oranges) signify that the latter profile was more highly conserved, and values greater than zero (blues to black) signify that the native residue was less conserved in the latter profile. Not along the diagonal, values less than zero indicate that exchangeability of the native residue to the indicated residue along the y axis was higher in the latter profile, whereas values greater than zero indicate that exchangeability was lower in the latter profile.

**S3 Fig. Comparison of individual exchangeability rates.** (A) Scatterplots of each exchangeability rate as observed in a PSI-BLAST profile compared to design profile. Both the x and y axes represent the exchangeability frequency of a native amino acid to a specific, non-native amino acid, with PSI-BLAST exchangeability rates along the x axis, and design exchangeability rates along the y axis. For reference, a grey line drawn is drawn along where the exchangeability rates would be equal between a PSI-BLAST and design profile. Both an adjusted *r*^2^and τ_*β*_ value is provided, along with the associated two-sided *p*-value. Lighter points are found along the linear regression model, and darker points represent outliers. (B) Measures of influence for individual exchangeability rate. The index listed along the x axis refers to each exchangeability rate, indexed in order alphabetically. For reference, in the first nineteen indices, the first index refers to the A to C mutation frequency, followed by the next eighteen indices that correspond with A to D through Y mutation frequencies. The measure of influential observation, or DFBETA index, is represented along the y axis. The height and direction of each bar corresponds with the change in regression model correlation coefficient without that particular observation. Influential outliers that have 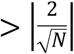 index value, or ± 0.106 threshold, are colored in black and are labeled with the associated mutation.

**S4 Fig. Correlation of dihedral angle RMSD and Cβ−Cβ distance deviation.** (A) The x-axis represents dihedral RMSD, measured in radians, and the y-axis represents contact proximity deviation, measured in Å. The hex bins shaded in grey are the number of residues within the deposited PDB structure have have both a C_β_−C_β_ distance deviation and dihedral angle RMSD within a bin. (B) Axes represent same metrices as in Panel A, normalized by z-score.

